# Italian norms and naming latencies for 357 high quality color images

**DOI:** 10.1101/492256

**Authors:** Eduardo Navarrete, Giorgio Arcara, Sara Mondini, Barbara Penolazzi

## Abstract

In the domain of cognitive studies on the lexico-semantic representational system, one of the most important means of ensuring well-suited experimental designs is using ecological stimulus sets accompanied by normative data on the most relevant variables affecting the processing of their items. In the context of image sets, color photographs are particularly suited for this aim as they reduce the difficulty of visual decoding processes that may emerge with traditional image sets of line drawings, especially in clinical populations. We provide Italian norms for a set of 357 high quality image-items belonging to 23 semantic subcategories. Data from several variables affecting image processing: age of acquisition, familiarity, lexical frequency, manipulability, name agreement, typicality and visual complexity; were collected from a sample of 255 Italian-speaking participants. Lexical frequency data were derived from the CoLFIS corpus. Furthermore, we collected data with on image naming latencies aimed at exploring how much of the variance in these latencies could be explained by the above mentioned critical variables. Multiple regression analyses on the naming latencies show classical psycholinguistic phenomena, such as the effects of age of acquisition and name agreement. In addition, manipulability is also a significant predictor. The described Italian normative data and naming latencies are available for download as supplementary material.

## Introduction

Object naming is perhaps the most widely exploited task for studying lexical access during speech production. Decades of research using this paradigm have allowed to identify some of the variables that influence the speed and the accuracy with which words are retrieved from the mental lexicon. It is undeniable that this advance in psycholinguistic knowledge of speech production processes has been closely linked to the appearance of standardized sets of stimuli to be named.

In this respect, one of the first and most influential normative data sets is the battery of Snodgrass and Vanderwart [1]. This set consists of 260 black and white line drawings containing values for four relevant variables affecting cognitive processing during object naming: familiarity, image agreement, name agreement and visual complexity (see for a color version of the battery, [2]). Normative data studies have recently started to use a more ecological type of stimuli, where line-drawings are being replaced by photographs. Under the assumption that photographs provide more surface and texture details than line-drawings, it has been hypothesized that photographs would accelerate visual processing and that this, in turn, could accelerate the lexicalization process. Congruent with this hypothesis, Salmon, Mateshon and McMullen [3] showed, for instance, that photographs of tools are named faster than their corresponding line-drawings. Salmon and colleagues interpreted these results as congruent with the notion that the automatic activation of the motor cortex areas associated with the use of tools (e.g., [4]) is facilitated by photographic stimuli in comparison to line-drawing stimuli. These results indicate the importance of controlling for visual features of the items to be named, at least for the semantic category of tools. At the same time, visual features are likely to play an important role in other semantic categories beyond that of tool [5]. Perceptual characteristics of items also influence other cognitive domains besides word production. For instance, recent memory studies reported that the perceptual characteristics of the items to be learned constitute a cue that impacts memory predictions for those items. Specifically, items that are easier to perceive during encoding generate higher judgments of learning (a.k.a. JOLS), despite the fact that ease of perception does not generally influence subsequent recall performance. Such a phenomenon is observed with word stimuli [6] as well as with picture stimuli [7, 8].

In addition, color is an essential attribute of objects and therefore provides a more realistic representation. The greater richness provided by color photographs compared to black and white photographs has been shown to improve object perception (e.g., [9,10]), although it does not seem to ameliorate semantic processing [11]. Consequently, normative studies have started to use more ecological color photographs instead of black and white stimuli (e.g., [12–14]).

Normative data studies have not only been concerned about the representation modality of the stimuli (black and white line-drawings or color photographs), but also about the number of critical variables included in these studies as predictors of lexical access during speech production. Compelling evidence shows that, apart from the four variables presented in the original study by Snodgrass and Vanderwart [1], other variables affect the lexicalization process, like, for instance, the age at which a word is first learned. Specifically, early acquired words tend to be named faster and more accurately than late acquired words. This phenomenon, known as the Age of Acquisition effect (AoA), is not exclusive to object naming but is widespread across several lexico-semantic processes such as semantic categorization [15], reading [16], or the probability of retrieving a word from the mental lexicon during language production [17], (for a review, see [18]). Another relevant variable determining the speed and accuracy with which words are retrieved is word frequency. In object naming tasks, high frequency words are named faster and more accurately than low frequency words [19, 20]. Critically, such an advantage is absent when the task does not require lexical access, as for instance when participants are asked to indicate whether previously presented words denoted the objects depicted in target pictures (i.e., old/new decision task; [21]). This evidence suggests that the phenomenon is mainly ascribed to a lexical-phonological level of processing [22]. A third critical predictor for object naming is manipulability. Broadly speaking, manipulability refers to the possibility of manual interactions with a specific object. It has been reported recently that items with a high level of manipulability are named faster than items with a low level of manipulability [3, 23]. Although manipulability remains a vague variable (see for discussion, [13]), and the level of processing at which the advantage takes place is still unclear (see for discussion, [23]), the phenomenon has been replicated in different languages.

In sum, since the original study of Snodgrass and Vanderwart [1], researchers have focused on more ecological stimuli such as color photographs and, at the same time, have increased the number of standardized variables crucially affecting the performance in lexical tasks. The objective of the present study is to offer researchers working with Italian-speaking participants a standardized set of 357 high quality color photographs ascribable to a high number of subcategories, along with norms for eight variables affecting image processing: AoA, familiarity, lexical frequency, manipulability, two name agreement measures (see below), typicality and visual complexity. To this end, we standardized in the Italian language the set of images provided by Moreno-Martínez and Montoro [24]. In addition to the cross-linguistic validation, as a novelty, we conducted an oral naming study in order to identify the more relevant predictors of naming latencies for the current set of images.

## Method

### Participants

A total of 255 healthy Italian native speakers (198 females, 57 males; mean age ± 21.29; sd. ± 3.54; 238 right-handed, 17 left-handed) participated in the rating study. In addition, twenty native Italian speakers (15 females, 5 males; mean age ± 20.6; sd. ± 1.39; 19 right-handed, 1 left-handed) took part in the naming study. All 275 participants had normal or corrected-to-normal vision and were students from Padova University, who attended a degree course in Psychology and participated to obtain credits.

*Ethical statement*. The procedures have been approved by the Ethical Committee for Psychological Research of the University of Padova before the study began (Protocol number: 1395-CB7EFAF01EE7652929D155AFEE6552FF; Title: *Mechanisms of Word Retrieval in Spoken Language Production*). Additionally, participation was voluntary and participants were explained that they were free to suspend their participation in the experiments at any time and for any cause.

### Stimuli

Stimuli were the freely available set of color photographs by Moreno-Martínez and Montoro [24]. This set is composed of 360 high-quality color photographs belonging to 23 semantic subcategories. Specifically, ten subcategories were selected from the living domain: animals, birds, body parts, dried fruits, insects, flowers, fruits, sea creatures, trees, vegetables; twelve subcategories were selected from the nonliving domain: buildings, clothing, foodstuff, furniture, jewelry, kitchen utensils, musical instruments, office material, sports/games, tools, vehicles, weapons; and finally, nonliving natural things (e.g., containing items like ‘mountain’, ‘stone’, etc.). Because of three items of the original database (i.e., “zarajo” and “porra” from the foodstuff category and “churrera” from kitchen utensils) were idiosyncratic items of Spanish culture and were impossible to translate into Italian, they were not included in the Italian set of images. Therefore, we collected norms for a total of 357 items. As described by Moreno-Martínez and Montoro [24], the color photographs were taken by the authors, the images were then modified to remove their original backgrounds (except for the nonliving natural things) and placed on a plain white background. Images have a mean dimension of 265×223 pixels and, for each category susceptible to being oriented, half of the items were left-facing and the other half were right-facing. Some examples of items are presented in Figure 1.

**Fig 1.**
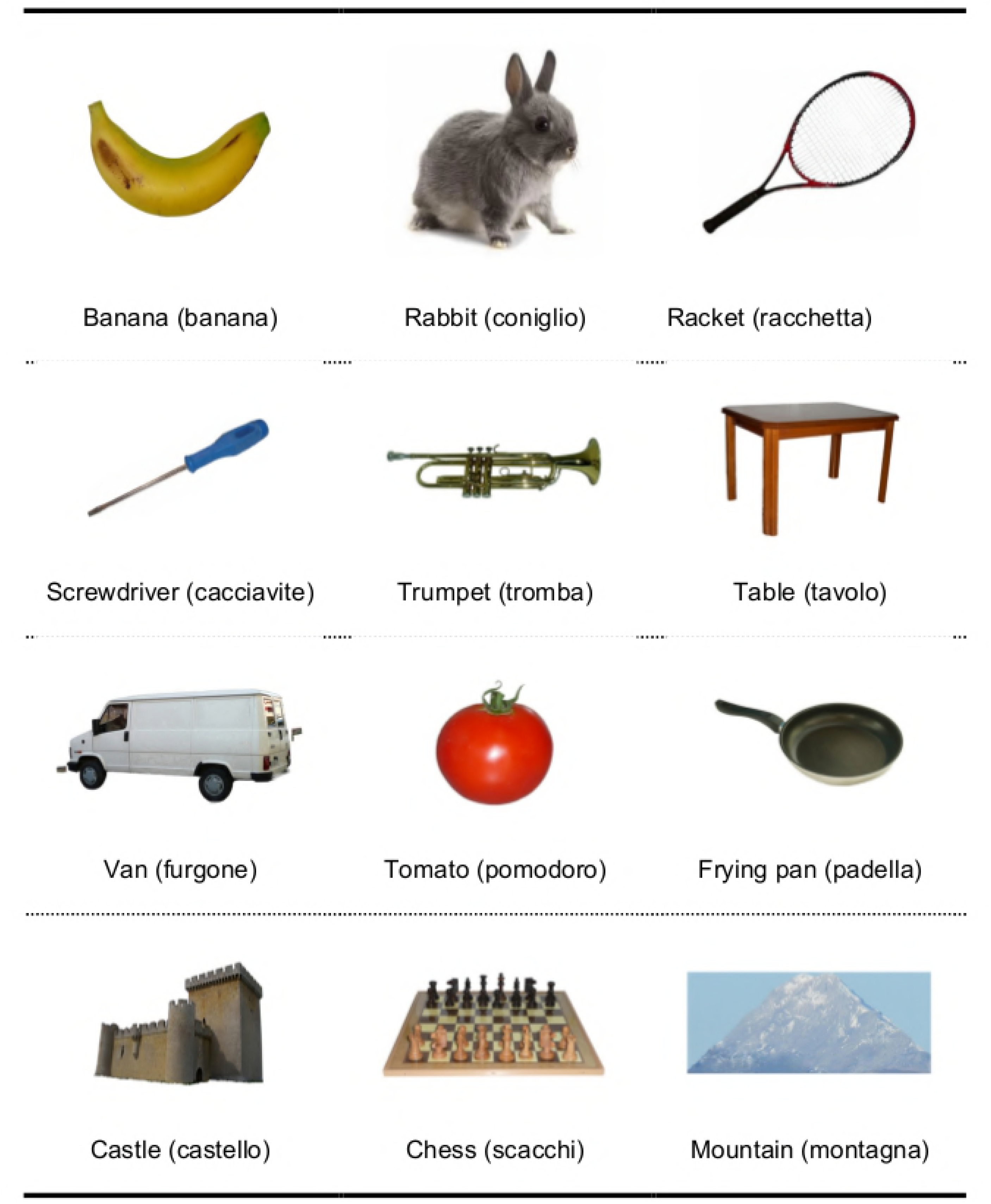
Some examples of the set of color photographs. Italian names are given in brackets.

### Procedure – Rating tasks

To guarantee uniformity across studies, the experimental procedure was kept as similar as possible to that used in Moreno-Martínez and Montoro’s study [24]. Images were shown to a sample of 255 participants, who had to first typewrite on the computer keyboard the name of the object represented in each figure without any time pressure. Subsequently, participants were required to rate the five psycholinguistic variables included in the study: visual complexity, AoA, familiarity, manipulability and typicality. Visual complexity was always the first rating assessed, and typicality was always the last. The other three ratings (AoA, familiarity, manipulability) were randomly presented across subjects. To guarantee a high consistency with the original study, the 357 images to be evaluated were divided into three lists (A, B, C) containing a similar number of exemplars from each of the 23 subcategories and a total of 119 items each. Participants were randomly assigned to perform the task on one of the three lists into which the entire image set was divided. Specifically, 81 were assigned to list A (64 females and 17 males; age: 21.46±2.61), 90 were assigned to list B (69 females and 21 males; age: 21.47±5.24) and 84 were assigned to list C (65 females and 19 males; age: 20.94±1.84). Participants performed the task individually in one session lasting about 90 minutes (with self-administered rest periods), and were tested simultaneously in groups of about 30 people in the same room.

The task was preceded by a practice phase in which participants were required to typewrite the name and subsequently rate 10 items not belonging to the main dataset for the same described variables. This practice phase was aimed at enabling participants to become familiar with the task and to develop anchor points useful for rating the subsequent stimulus material. Image delivery and participants’ responses were controlled by E-Prime 2.0 software (Psychology Software Tools, Inc., Pittsburgh, PA). The pictures were displayed on computers with Dell monitor 21.52''; participants’ distance from the screen was approximately 60 cm. Each image was preceded by a 500 ms-fixation cross, and remained visible for 3000 ms for the naming task or until a response was given for the rating task.

In the *Typewriting* (naming) task participants were asked to write the object represented in each image, trying to give specific names rather than general names indicating the category they belong to. In other words, participants were encouraged to provide the specific name for each image and avoid general or categorical names (e.g., “rose”, instead of general names like “flower” or “plant”). Participants typed the name of each image on the computer keyboard without time pressure. Participants were also instructed to type the initials NC for “don’t know” (NC = “non conosco” in Italian) if they did not recognize the object in the image and to type PL for “tip of the tongue” (PL = “punta della lingua” in Italian) if they knew what the object in the image represented but they were momentarily unable to remember its name. They had to type NR for “don’t remember” (NR = “Non ricordo” in Italian) if they recognized the object in the image but did not know if there was a word to name it. Typed responses were saved by the program. Name agreement measure was calculated based on the percentage of participants who named the item according to its canonical name. Two measures of name agreement were calculated: the percentage of participants who gave the canonical/dominant name to each specific item and the H statistic. The H statistic is a logarithmic function describing the different names that an item received and the proportion of participants giving each name [1], capturing information about the dispersion of the names. It has been shown that the H statistic captures more information about the variability of names across participants than does the simple percentage-of-agreement measure [1, 25, 26]. The H statistic is calculated as follows:

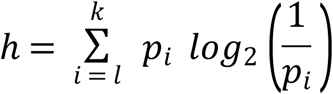

where k refers to the number of different names given to each item, and *p_i_* is the value for each name as a proportion of all the alternative names. For example, if only one name is given to a photo, H is zero; if two names occur with equal frequencies, H is 1. Thus, H increases with the number of given responses for the same item, and it is higher if the alternatives have similar probabilities.

*Rating tasks*. The rating tasks were performed by pressing numbers on the keyboard that corresponded to their evaluation. In line with the original study [24], in the visual complexity ratings, participants were required to “rate the visual complexity of the image itself, rather than that of the object it represents”, evaluating “the amount of details, intricacy of lines, pattern and quantity of colors presented in the image”’ using a 5-point scale (1 = very simple, 5 = very complex). To evaluate the remaining variables (AoA, familiarity, manipulability and typicality), participants were asked to “rate the object represented rather than the image itself”. For the AoA, familiarity, manipulability and typicality ratings, the image was presented together with the canonical name of the item (i.e., the expected name). Additionally, in the typicality rating task, the category name of the item was also provided (e.g., “fruit”, for the item “lemon”).

For AoA participants were instructed to rate the age at which they thought they had first learned each word using a seven-point scale (1 = 0-2 years, 7 = 13 years or more). In the familiarity rating task, participants were instructed to rate each item by assessing how often they thought they were exposed to each of them on the basis of how frequently they thought about the concept, and how frequently they came into contact with the concept (both directly through real-life exemplars and in a mediated way, as represented in the media), using a 5-point scale (1= very unfamiliar, 5= very familiar). In the manipulability rating task, participants were instructed to rate each item by assessing “the degree to which using a human hand is necessary for this object to perform its function”, using a 5-point scale (1 = never necessary, 5 = totally indispensable). The typicality test aims to measure the degree to which a concept is a representative exemplar of its category. Participants scored how representative of its category they thought an exemplar was (e.g., “ship” for “vehicle”) using a 5-point scale (1= not at all prototypical, 5= very prototypical). Lexical frequency values were retrieved from the CoLFIS database (which comprises 3,798,275 lexical occurrences; [27]).

### Procedure – Oral naming task

In the oral naming task, participants were asked to name the photos as fast and as accurately as possible. They were instructed to remain silent in case they did not recognize or did not know the name of the object. The pictures were displayed on computers Dell OptiPlex 520, Pentium 4 3.0 GHz, with Dell monitor 21.52''. Participants were seated approximately 50 cm from the computer screen, wearing a headset microphone. Naming latencies were measured from the appearance of the stimulus. An experimental trial contained the following events: (1) a fixation cross in the center of the screen for 500 ms; (2) a blank screen for 500 ms; (3) and the target stimulus for 3000 ms or until the participant responded. The next trial began 1500 ms after the onset of participants’ response or the disappearance of the stimulus. Stimulus presentation and response recording were controlled by DMDX program [28]. The set of items was presented in a different random order for each participant. Due to a mistake during the randomization of the lists, one item (i.e., Square.jpg) was not presented. There was a short pause after 50 trials. The first two trials at the beginning of the experiment and after each pause were warm-up trials containing 2 filler images not presented in the experimental set.

## Results – Normative study

### Descriptive Results

Appendix S1 provides descriptive statistics (means and standard deviations) of each variable assessed in the rating tasks (AoA, familiarity, manipulability, typicality, visual complexity) for all of the dataset stimuli (grouped by semantic category). In line with the original Spanish dataset, word-form frequency of the dominant name is expressed as a natural logarithm. Two measures of name agreement were provided in the Appendix S1. In Appendix S2 we reported the alternative names each item received; while in Appendix S3 we reported indexes of individual item analysis, including an item difficulty measure and two indexes of item discrimination based on item-test correlations (biserial and point-biserial). Table 1 presents summary statistics for all the variables, and Table 2 the same statistics separately for all the subcategories. Table 3 shows Pearson’s correlations among the variables.

**Table 1.** Summary statistics for all the variables.

|  | Agr-P | Agr-H | Freq | AoA | Vis Com | Fam | Man | Typ |
| --- | --- | --- | --- | --- | --- | --- | --- | --- |
| mean | 0.56 | 1.49 | 7.61 | 3.99 | 2.61 | 3.10 | 2.84 | 3.48 |
| sd | 0.35 | 1.01 | 2.18 | 1.36 | 0.71 | 0.99 | 1.26 | 0.83 |
| median | 0.64 | 1.41 | 7.73 | 3.89 | 2.52 | 3.03 | 2.95 | 3.56 |
| max | 1 | 3.51 | 13.23 | 6.83 | 4.43 | 4.93 | 4.96 | 4.85 |
| min | 0 | 0 | 0 | 1.42 | 1.17 | 1.02 | 1.03 | 1.44 |
| range | 1 | 3.51 | 13.23 | 5.41 | 3.26 | 3.91 | 3.93 | 3.41 |
| kurtosis | -1.44 | -1.24 | 1.07 | -0.94 | -0.61 | -1 | -1.43 | -0.70 |
| skewness | -0.28 | 0.14 | -0.51 | 0.14 | 0.25 | 0.12 | 0.07 | -0.37 |
| Q1 | 0.23 | 0.58 | 6.39 | 2.99 | 2.04 | 2.33 | 1.56 | 2.90 |
| Q3 | 0.90 | 2.39 | 8.98 | 5.18 | 3.12 | 3.93 | 3.92 | 4.16 |
| Mode | 0 | 0 | 7.02 | 5.63 | 1.82 | 2.6 | 1.15 | 4.16 |

Note. Agr-P = Percentage of name agreement; Agr-H= H statistic on name agreement; Freq = Lexical frequency (natural logarithm); AoA =Age of Acquisition; Vis Com = Visual Complexity; Fam = Familiarity; Man=Manipulability; Typ = Typicality.

**Table 2.** Summary statistics for all the variables for each category.

| Category | Agr-P |  | Agr-H |  | Freq |  | AoA |  | Vis Com |  | Fam |  | Man |  | Tip |  |
| --- | --- | --- | --- | --- | --- | --- | --- | --- | --- | --- | --- | --- | --- | --- | --- | --- |
|  | M | SD | M | SD | M | SD | M | SD | M | SD | M | SD | M | SD | M | SD |
| Clothes | 0.75 | 3.98 | 0.27 | 0.49 | 0.93 | 0.62 | 3.17 | 0.97 | 8.36 | 1.02 | 1.07 | 3.61 | 1.39 | 1.94 | 1.01 | 3.35 |
| Tress | 0.20 | 2.65 | 0.30 | 0.33 | 0.72 | 0.23 | 4.93 | 0.66 | 7.47 | 0.49 | 2.10 | 3.62 | 0.84 | 2.81 | 0.79 | 1.45 |
| Food | 0.37 | 3.18 | 0.28 | 0.58 | 0.71 | 0.52 | 4.42 | 0.95 | 6.58 | 0.71 | 2.29 | 3.08 | 1.47 | 2.51 | 1.99 | 3.42 |
| Animals | 0.79 | 2.56 | 0.25 | 0.27 | 1.02 | 0.42 | 3.46 | 0.73 | 7.34 | 0.81 | 0.92 | 3.61 | 1.25 | 3.11 | 1.56 | 1.39 |
| Weapons | 0.61 | 2.12 | 0.37 | 0.73 | 0.92 | 0.67 | 4.44 | 0.35 | 8.35 | 0.92 | 1.21 | 3.17 | 1.10 | 2.56 | 1.34 | 4.11 |
| Tools | 0.39 | 2.53 | 0.37 | 0.39 | 0.97 | 0.42 | 4.82 | 0.58 | 7.18 | 0.65 | 1.88 | 3.42 | 1.10 | 2.06 | 1.72 | 4.33 |
| Desk Material | 0.63 | 3.74 | 0.35 | 0.59 | 0.96 | 0.30 | 3.95 | 0.90 | 7.41 | 0.86 | 1.39 | 3.69 | 1.14 | 2.01 | 3.27 | 4.54 |
| Marine Creatures | 0.44 | 2.13 | 0.37 | 0.57 | 0.94 | 0.60 | 4.82 | 0.73 | 6.10 | 0.89 | 1.68 | 3.08 | 1.43 | 2.73 | 2.36 | 1.63 |
| Buildings | 0.54 | 2.79 | 0.32 | 0.68 | 0.89 | 0.36 | 4.33 | 0.89 | 9.10 | 1.02 | 1.83 | 2.91 | 1.36 | 3.41 | 2.09 | 1.99 |
| Flowers | 0.46 | 2.95 | 0.40 | 0.51 | 1.32 | 0.19 | 4.56 | 0.82 | 6.43 | 0.66 | 1.97 | 3.82 | 1.35 | 2.62 | 2.08 | 1.82 |
| Fruits | 0.56 | 3.46 | 0.42 | 0.42 | 1.09 | 0.31 | 3.64 | 1.02 | 7.09 | 0.91 | 1.28 | 3.62 | 1.57 | 1.89 | 2.54 | 3.39 |
| Nuts | 0.45 | 3.06 | 0.32 | 0.32 | 0.82 | 0.56 | 4.17 | 0.68 | 6.89 | 0.80 | 1.91 | 3.46 | 0.67 | 2.06 | 1.47 | 3.09 |
| Jewelry | 0.49 | 3.16 | 0.33 | 0.60 | 0.89 | 0.70 | 4.17 | 1.09 | 7.17 | 0.99 | 1.63 | 3.55 | 1.23 | 2.83 | 2.67 | 3.51 |
| Insects | 0.61 | 2.95 | 0.30 | 0.53 | 0.88 | 0.16 | 3.79 | 0.87 | 6.63 | 0.58 | 1.49 | 3.75 | 1.28 | 2.97 | 1.50 | 1.33 |
| Furniture | 0.67 | 4.10 | 0.37 | 0.49 | 1.25 | 0.66 | 3.39 | 0.83 | 8.36 | 0.88 | 1.21 | 3.72 | 1.22 | 2.48 | 2.01 | 2.84 |
| Nature | 0.62 | 3.47 | 0.34 | 0.71 | 1.03 | 0.44 | 3.08 | 0.88 | 9.90 | 0.89 | 1.40 | 3.34 | 1.06 | 2.67 | 1.64 | 1.44 |
| Body parts | 0.64 | 3.98 | 0.34 | 0.51 | 1.01 | 0.94 | 3.06 | 0.98 | 9.86 | 0.77 | 1.09 | 3.80 | 1.68 | 2.37 | 1.52 | 2.09 |
| Sports/Games | 0.57 | 3.10 | 0.34 | 0.76 | 0.94 | 0.94 | 3.55 | 0.63 | 6.76 | 0.71 | 1.32 | 3.39 | 1.21 | 2.82 | 2.27 | 3.93 |
| Musical instruments | 0.64 | 2.66 | 0.31 | 0.67 | 0.79 | 0.21 | 4.48 | 0.64 | 7.83 | 0.77 | 1.36 | 3.75 | 1.08 | 3.29 | 1.65 | 4.71 |
| Birds | 0.55 | 2.52 | 0.34 | 0.46 | 0.93 | 0.19 | 4.31 | 0.80 | 7.03 | 0.60 | 1.66 | 3.17 | 1.40 | 3.30 | 1.02 | 1.30 |
| Kitchen utensils | 0.44 | 3.59 | 0.39 | 0.61 | 1.22 | 0.26 | 4.39 | 1.07 | 6.14 | 0.93 | 1.78 | 3.60 | 1.53 | 2.22 | 3.07 | 4.31 |
| Vehicles | 0.62 | 3.48 | 0.34 | 0.58 | 0.85 | 0.77 | 3.48 | 0.81 | 9.16 | 1.10 | 1.30 | 3.25 | 0.88 | 3.11 | 1.91 | 3.61 |
| Vegetables | 0.60 | 3.42 | 0.39 | 0.47 | 1.08 | 0.27 | 4.15 | 0.84 | 7.35 | 0.62 | 1.38 | 3.57 | 1.14 | 2.08 | 1.36 | 3.30 |

Note. Agr-P = Percentage of name agreement; Agr-H= H statistic on name agreement; Freq = Lexical frequency (natural logarithm); AoA =Age of Acquisition; Vis Com = Visual Complexity; Fam = Familiarity; Man=Manipulability; Typ = Typicality.

**Table 3.**
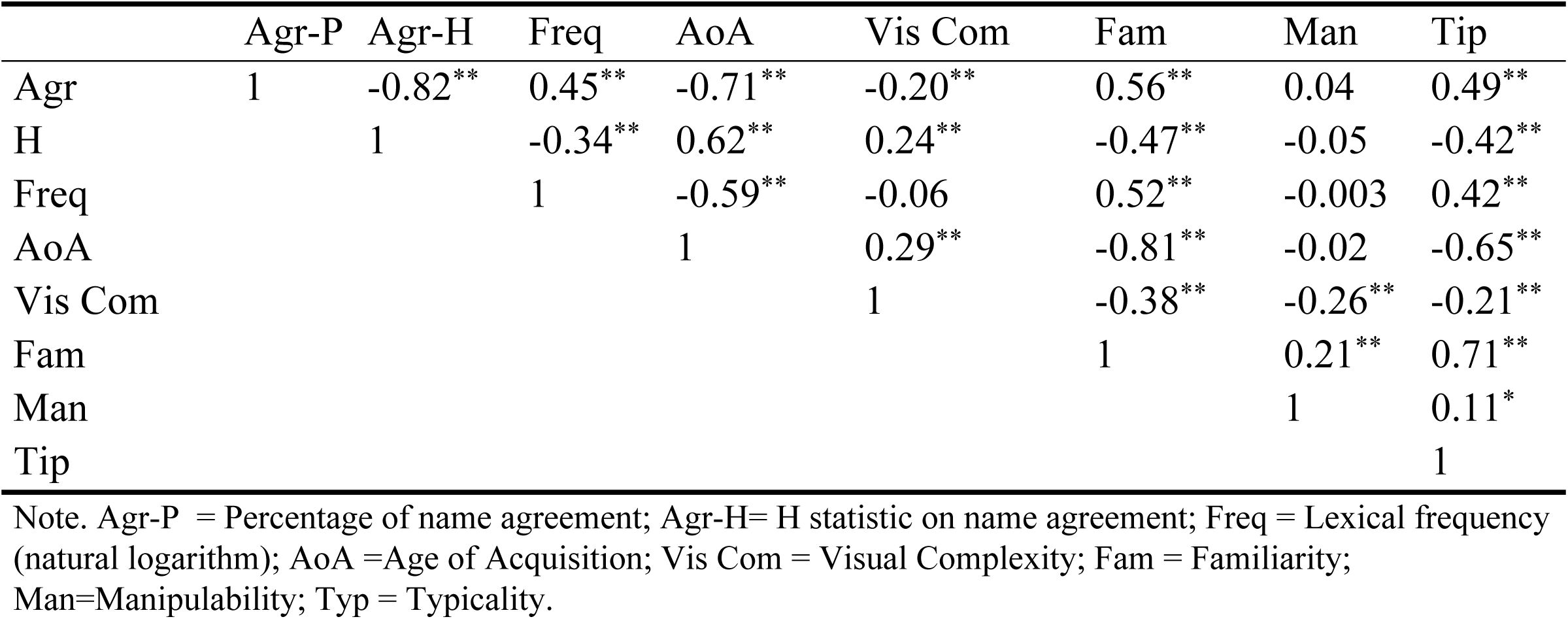
Correlation matrix among the variables.

|  | Agr-P | Agr-H | Freq | AoA | Vis Com | Fam | Man | Tip |
| --- | --- | --- | --- | --- | --- | --- | --- | --- |
| Agr | 1 | -0.82** | 0.45** | -0.71** | -0.20** | 0.56** | 0.04 | 0.49** |
| H |  | 1 | -0.34** | 0.62** | 0.24** | -0.47** | -0.05 | -0.42** |
| Freq |  |  | 1 | -0.59** | -0.06 | 0.52** | -0.003 | 0.42** |
| AoA |  |  |  | 1 | 0.29** | -0.81** | -0.02 | -0.65** |
| Vis Com |  |  |  |  | 1 | -0.38** | -0.26** | -0.21** |
| Fam |  |  |  |  |  | 1 | 0.21** | 0.71** |
| Man |  |  |  |  |  |  | 1 | 0.11* |
| Tip |  |  |  |  |  |  |  | 1 |

Note. Agr-P = Percentage of name agreement; Agr-H= H statistic on name agreement; Freq = Lexical frequency (natural logarithm); AoA =Age of Acquisition; Vis Com = Visual Complexity; Fam = Familiarity; Man=Manipulability; Typ = Typicality.

### Reliability

To determine the reliability of our data, we correlated the variables between the items sharing the same dominant name in the present study and other studies [24]. In particular, 357 items overlapped with Moreno-Martínez and Montoro [24], 50 with Adlington, Laws and Gale [29], 69 with Brodeur et al. [12], 113 with Moreno-Martínez, Montoro and Laws [30], 107 with Snodgrass & Vanderwart [1], 81 with the English version of Viggiano et al. [14], and 80 with the Italian version of Viggiano et al. [14]. Pearson’s correlations are reported in Table 4. Correlations fluctuated between .28 and .98.

**Table 4.** Correlations between current stimuli and those of Moreno-Martínez and Montoro (2012), Adlington et al. (2009), Brodeur et al. (2010), Moreno-Martínez et al. (2011), Snodgrass & Vanderwart (1980) and Viggiano et al. (2004).

|  | Items (n) | Agr-P | Freq | AoA | Vis Com | Fam | Man |
| --- | --- | --- | --- | --- | --- | --- | --- |
| Moreno-Martínez & Montoro | 357 | 0.65 | 0.57 | 0.87 | 0.94 | 0.88 | 0.96 |
| Adlington et al.'s | 50 | 0.53 | 0.77 | 0.85 | 0.64 | 0.76 | NA |
| Brodeur et al.'s | 69 | 0.38 | NA | NA | 0.66 | 0.65 | 0.52 |
| Moreno-Martínez et al | 113 | 0.75 | 0.62 | 0.9 | 0.88 | 0.87 | 0.98 |
| Snodgrass & Vanderwart | 107 | 0.36 | 0.58 | 0.79 | 0.75 | 0.85 | NA |
| Viggiano et al. (English) | 81 | 0.28 | NA | NA | 0.79 | 0.74 | NA |
| Viggiano et al. (Italian) | 80 | 0.31 | NA | NA | 0.84 | 0.82 | NA |

Note. Agr-P = Percentage of name agreement; Agr-H= H statistic on name agreement; Freq = Lexical frequency (natural logarithm); AoA =Age of Acquisition; Vis Com = Visual Complexity; Fam = Familiarity; Man=Manipulability; Typ = Typicality.

## Results – Oral naming study

### Data analysis

Naming latencies and accuracy were determined offline using the CheckVocal software [30]. Responses in which participants produced an incorrect name (e.g., semantic superordinates such as “tool” instead of “pliers”; semantic coordinates such as “boat” instead of “ship”), missing responses and verbal dysfluencies were excluded from the analysis. Following those criteria a total of 30.2% of the data were excluded (15.2% of error response; 15% of no-responses and 0.03% of voice key problems). Fourteen items did not elicit correct responses and were excluded from the analysis. Correct responses have a mean latency of 1203 ms with a standard deviation of 361 ms. In a first descriptive level of analysis, we performed correlations between naming latencies and the eight variables of the normative study.

In a second type of analysis, we assessed how much of the variance in the naming latencies was explained by each of the above variables. Analysis was performed on the average of the naming latencies of each stimulus. In order to avoid problems of collinearity, we adopted the following approach: first, we assessed the correlations among the variables through a hierarchical clustering analysis using the *varclus* function of the “Hmisc” package [32] with the R statistical software [33]. This allowed us to identify clusters of variables (i.e., variables with a Spearman similarity coefficient > .35). This kind of analysis separates variables into clusters that can be scored as a single variable, thus resulting in data reduction. In a second step, we selected the more important variable within each cluster through likelihood ratio tests between the models containing separately each one of the identified variables in each cluster. Finally, a multiple regression analysis was used to explore how much of the variance in the naming latencies was explained by the variables. This second kind of analysis was performed on a subset of items that received on the normative typewriting task a name agreement value equal to or above 50% (i.e., items that elicited the expected name from at least half of the participants). This criterion was selected in order to exclude spurious influences, as for instance poor visual structural descriptions of the photos, or the impact of idiosyncratic linguistic characteristics of the target words in Italian ([34]; for an example of the influence of name agreement on picture naming latencies see [35]). Following this criterion, the analysis was performed on 196 items (see Appendix S4 for a further analysis including all items and mixed effect modeling).

### Results

As can be seen in Table 5, naming latencies correlated positively with H statistic, AoA and visual complexity and negatively with name agreement, lexical frequency, familiarity and typicality. No significant correlation was obtained between naming latencies and manipulability.

**Table 5.** Correlation between the average of naming latencies and the variables.

|  | Agr-P | Agr-H | Freq | AoA | Vis Com | Fam | Man | Typ | Df |
| --- | --- | --- | --- | --- | --- | --- | --- | --- | --- |
| Set A | -0.56** | 0.63** | -0.35** | 0.57** | 0.14** | -0.45** | 0.02 | -0.54** | 341 |
| Set B | -0.54** | 0.62** | -0.32** | 0.66** | 0.24** | -0.48** | 0.01 | -0.50** | 194 |

Note. Set A = 343 items; Set B = 196 items (items with agreement values in the normative study equal or above 0.5; see main text for details). See the Note in Table 1. Df = degrees of freedom. \*\* $p < .01$

Figure 2 shows the hierarchical clustering structure among the variables. Two clusters of highly correlated variables emerged. Agreement and H statistic formed a cluster and typicality, AoA and familiarity formed another one. The likelihood ratio test indicated that H statistic produced a significant increase in the explained variance compared to agreement (χ^2^ = 27.75, *p* < .001). In the second cluster, the likelihood ratio tests indicated that AoA produced a significant increase in the explained variance compared to typicality (χ^2^ = 58.47, *p* < .001) and familiarity (χ^2^ = 63.41, *p* < .001). Thus, the multiple regression analysis was conducted with five variables: H statistic, AoA, frequency, manipulability and visual complexity. Partial effects of the model are illustrated in Figure 3. As can be seen in Table 6, H statistic, AoA and manipulability were significant predictors of naming latencies (R^2^= 0.611). Specifically, faster naming latencies were obtained for items with higher H statistic values, acquired early in life and with higher manipulability rating. No effects of visual complexity and lexical frequency were obtained (see Supplemental material for similar findings using a different type of analysis). Tolerance statistics were all above .5 and the average of the variance inflation factor (VIF) was 1.38, suggesting that collinearity was not a problem of the regression model [36].

**Fig 2.**
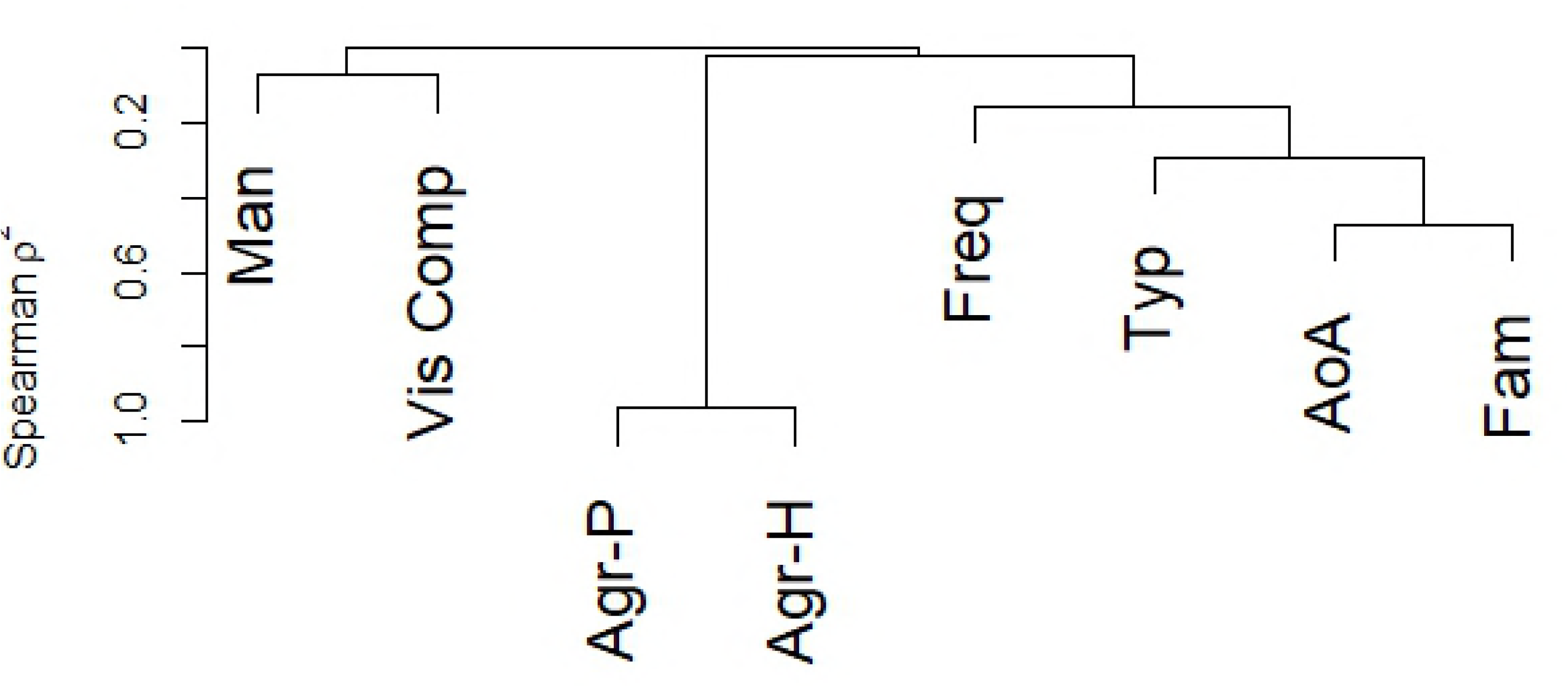
Hierarchical clustering analysis using Spearman’s p^2^ for the eight explored variables. See the Note in Table 1.

**Fig 3.**
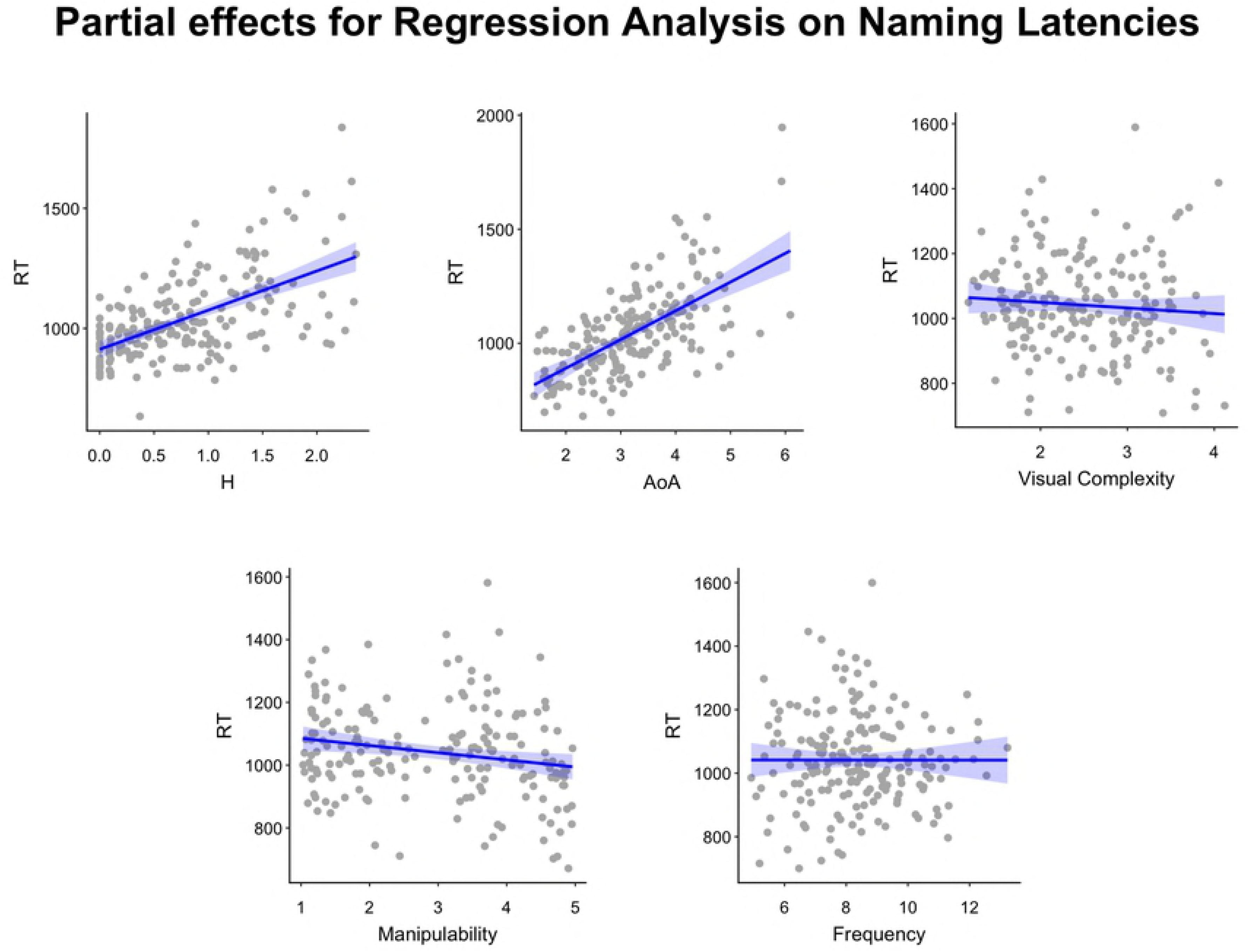
The figure shows the partial effects for the regression model on Naming Latencies. Each panel represents the partial effect of a predictor included in the regression. The blue lines represent the regression for that partial effect. The shaded blue represents the area delimiting the standard error of the estimate. The gray points indicate the partial residuals associated with the predictor.

**Table 6.** Results of the multiple regression analysis, with naming latencies as the criterion variable and H statistic, age of acquisition, visual complexity, manipulability and lexical frequency as predictors variables.

|  | Estimate | St.Error | t value | Pr<br>(>\|t\|) | Tolerance | VIF |
| --- | --- | --- | --- | --- | --- | --- |
| Intercept | 620.952 | 93.067 | 6.672 | < .001 |  | - |
| H statistic | 163.114 | 18.523 | 8.806 | < .001 | 0.82 | 1.21 |
| Age of acquisition | 125.974 | 14.729 | 8.552 | < .001 | 0.54 | 1.84 |
| Visual complexity | -17.335 | 17.115 | -1.013 | 0.312 | 0.75 | 1.32 |
| Manipulability | -22.860 | 8.740 | -2.615 | 0.009 | 0.85 | 1.16 |
| Lexical frequency | -0.064 | 7.382 | -0.009 | 0.993 | 0.71 | 1.39 |

Notes: VIF = variance inflation factor. Degrees of freedom = 190.

## Discussion

The present study provides Italian norms for 357 high quality color images from the set of Moreno-Martínez and Montoro [24]. Several psycholinguistic variables that have been showed to affect latency and accuracy during object naming are included: agreement, H statistic, word-lexical frequency, age of acquisition (AoA), visual complexity, familiarity, manipulability and typicality. As reported in other normative studies (e.g., [37, 38]), the variables are highly correlated. Critically, the correlation pattern we obtained matches to a large extent the one reported in the original Moreno-Martínez and Montoro’s study in Spanish.

A second aim of this study, which is the most original part in the Italian validation of Moreno-Martínez and Montoro’s database, was to explore how much of the variance in the latencies of the oral naming task could be explained by the above mentioned critical variables. A first descriptive analysis showed that all the variables, except manipulability, correlated with naming latencies. In a second analysis, in order to reduce multicollinearity problems, we separated the variables into clusters through a hierarchical cluster analysis and then we selected the most relevant variable for each cluster. The regression analysis using the five remaining variables as predictors showed a significant effect of H statistic, AoA and manipulability. Specifically, the items that tended to elicit a similar name from participants in the typewriting naming normative study (i.e., lower H index) were named faster (e.g., [35]). At the same time, items acquired early in life were named faster than items acquired late in life, replicating the well-known phenomenon of age of acquisition [18]. In addition, manipulability also modulated naming latencies once the control variables of name agreement (i.e., H statistic), age of acquisition and visual complexity were taken into account. Specifically, items that were rated as more manipulable in the rating study were named faster in the oral naming task. This result suggests that manipulability is a critical variable affecting speech production in picture naming, replicating previous recent findings [13, 23, 39]. An apparently unexpected outcome could be the lack of a significant effect of lexical frequency, since traditionally this variable has been demonstrated to be a very reliable predictor of naming latencies [19, 22]. However, the lack of lexical frequency is congruent with recent findings in naming tasks showing no lexical frequency effects if AoA has been also included in the statistical analysis as a predictor. This suggests that AoA is a more reliable predictor, which assimilates part of the effect tied to frequency [17, 40].

To our knowledge, this is the first study to provide such a high number of psycholinguistic indexes for such a high number of quality color images in Italian. Examining all these variables in detail is of critical relevance in object naming research, as well as in other cognitive research domains, such as memory or object perception. Having well-controlled and ecological stimulus sets is also of critical relevance in clinical and neuropsychological domains [41], both to improve assessment procedures, and to disclose which processing level can be most impaired in patients’ elaboration failures [11]. This normative work may help item selection for the design of experimental work and clinical trials.

## Acknowledgments

Part of this research was funded by grants awarded by the University of Padua to S.M. and B.P. The authors report no conflicts of interest associated with this study and state there has been no significant financial support that could have influenced its outcome. We thank Maria Merigo, Valeria Baldan and Lorenzo Bragato for their help with data collection.

## Supporting Information

**S1 Appendix. Normative psycholinguistic ratings for each item**.

**S2 Appendix. Alternative names and responses**.

**S3 Appendix. Indexes of individual item analysis**.

**S4 Appendix. Supplemental analysis including all items and mixed effect modeling**.

**S5 Appendix. Data file with normative psycholinguistic ratings and naming latencies for each item**.

**S1 Fig.**
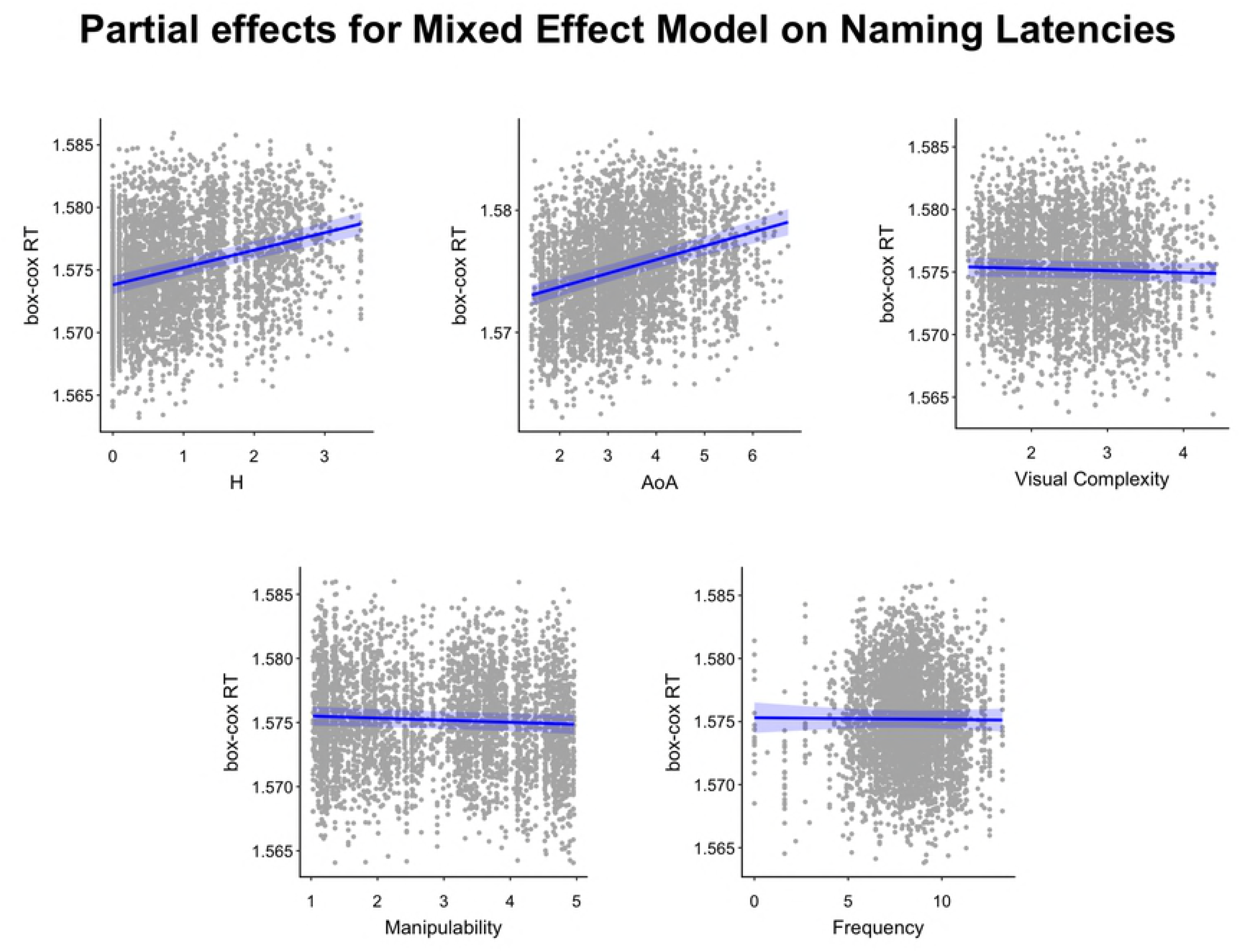

